# Classification of Multiple Emotional States from Facial Expressions in Mice using a Deep Learning-Based Image Analysis

**DOI:** 10.1101/2023.02.17.528923

**Authors:** Yudai Takana, Takuto Nakata, Hiroshi Hibino, Masaaki Nishiyama, Daisuke Ino

## Abstract

Facial expressions are widely recognized as universal indicators of underlying internal states in most species of animals, thereby presenting as a non-invasive measure for predicting physical and mental conditions. Despite the advancement of artificial intelligence-assisted tools for automated analysis of voluminous facial expression data in human subjects, the corresponding tools for mice still remain limited so far. Considering that mice are the most prevalent model animals for studying human health and diseases, a comprehensive characterization of emotion-dependent patterns of facial expressions in mice could extend our knowledge on the basis of emotions and the related disorders. Here, we present a framework for the development of a deep learning-powered tool for classifying mouse facial expressions. We demonstrate that our machine vision was capable of accurately classifying three different emotional states from mouse facial images. Moreover, we objectively determined how our classifier characterized the differences among the facial images through the use of an interpretation technique called Gradient-weighted Class Activation Mapping. Our approach is likely to facilitate the non-invasive decoding of a variety of emotions from facial images in mice.

## INTRODUCTION

Most species in the animal kingdom are proposed to use a facial expression as a nonverbal mean to externally display emotions, as described in a literature by Charles Darwin in the 19th century (1). To date, facial expressions have been well characterized in humans. Early studies by Paul Ekman (2) indicated the existence of the key facial expressions —happiness, sadness, anger, fear, surprise, disgust and neutral—, which can be discriminated based on distinct facial features. Through the combination of these key expressions, we humans are capable of conveying complex and diverse emotions to others during social interactions in daily life. In addition, recognizable changes in facial features appear in patients with disorders such as psychiatric symptoms, facial paralyses, and craniofacial anomalies (3–6). Therefore, facial expressions are of interest to both basic and clinical researchers as a non-invasive method for diagnosing physical and/or mental conditions.

Then, what about in non-human animals? Various species, such as primates, canines, and rodents, exhibit facial expressions in response to emotional events (7–12); therefore, the existence of facial expressions is likely in non-human animals. Considering that mice are one of the most commonly-used model animals for studying human health and disease, a quantitative assessment of facial expression in mice could advance our knowledge on emotions and related disorders. So far, a manual procedure and an automated tool for assessing pain from facial images of mice have been developed (9, 13), but these approaches do not afford us to analyze other emotional states. Recently, Dolensek and colleagues have reported a framework for a machine vision-assisted classification of multiple emotions from mouse facial images (11), but the procedure is unlikely applicable to multiple animals because the classifier, which utilized a traditional machine learning algorithm, was only able to learn features obtained from a single mouse. In the case of other species, including human (5, 14) and primates (15), machine vision-assisted approaches, particularly those utilizing deep learning (DL)-mediated image analyses, have achieved automated facial recognition with a high accuracy using large datasets derived from multiple individuals. Therefore, if advanced tools like these were available for the classification of multiple facial expressions in mice, it would be possible to extend the analyses to a diverse range of experiments.

In this study, we present a DL-based technique for categorizing multiple emotions from facial images of mice. By using an optimized facial videography technique for head-fixed animals, which affords us to clearly capture the dynamics of key facial parts such as ear, eye, and mouth in a single focus, we efficiently acquired a dataset consisting of thousands of mouse facial images for three emotional states: neutral, painful, and tickling. After subjecting the raw images in the dataset to preprocessing such as background masking and image resizing, we devised a convolutional neural network (CNN)-based classifier for the three emotional states using SqueezeNet (16), a deep CNN architecture with high accuracy and lower computational costs. Our model exhibited remarkable accuracy, registering a score of approximately 80%. Furthermore, by combining the analysis using Gradient-weighted Class Activation Mapping (Grad-CAM) (17), a technique producing the focus of CNN-based machine visions, with subsequent manual image analysis, we identified the potential key features for the image classification. Overall, our methodology presented here offers a streamlined approach for classifying a diverse range of mouse facial expressions.

## MATERIALS AND METHODS

### Animal surgery

All animal procedures were conducted following the guidelines of Kanazawa University and Osaka University. C57BL/6J female mice aged 2-4 months were obtained from CLEA Japan (Tokyo, Japan). Surgery to attach a stainless headplate (CP-1, Narishige, Tokyo, Japan) with a spacer (Fig. 1a) on the skull of the mouse using dental cement (Ketac Cem Easymix, 3M, Maplewood, MN, USA) was conducted under anesthesia of isoflurane. The depth of anesthesia was evaluated using the tail pinch method. Body temperature was maintained using a heating pad. To minimize inflammatory responses, carprofen (5 mg/kg; i.p.), a non-steroidal anti-inflammatory drug, and buprenorphine (0.1 mg/kg, i.p.), an opioid analgesic, were administered after the surgery. Mice were returned to cages, and were raised in cages under a regular 12-h dark/light cycle with ab libitum access to food and water.

**Fig. 1.**
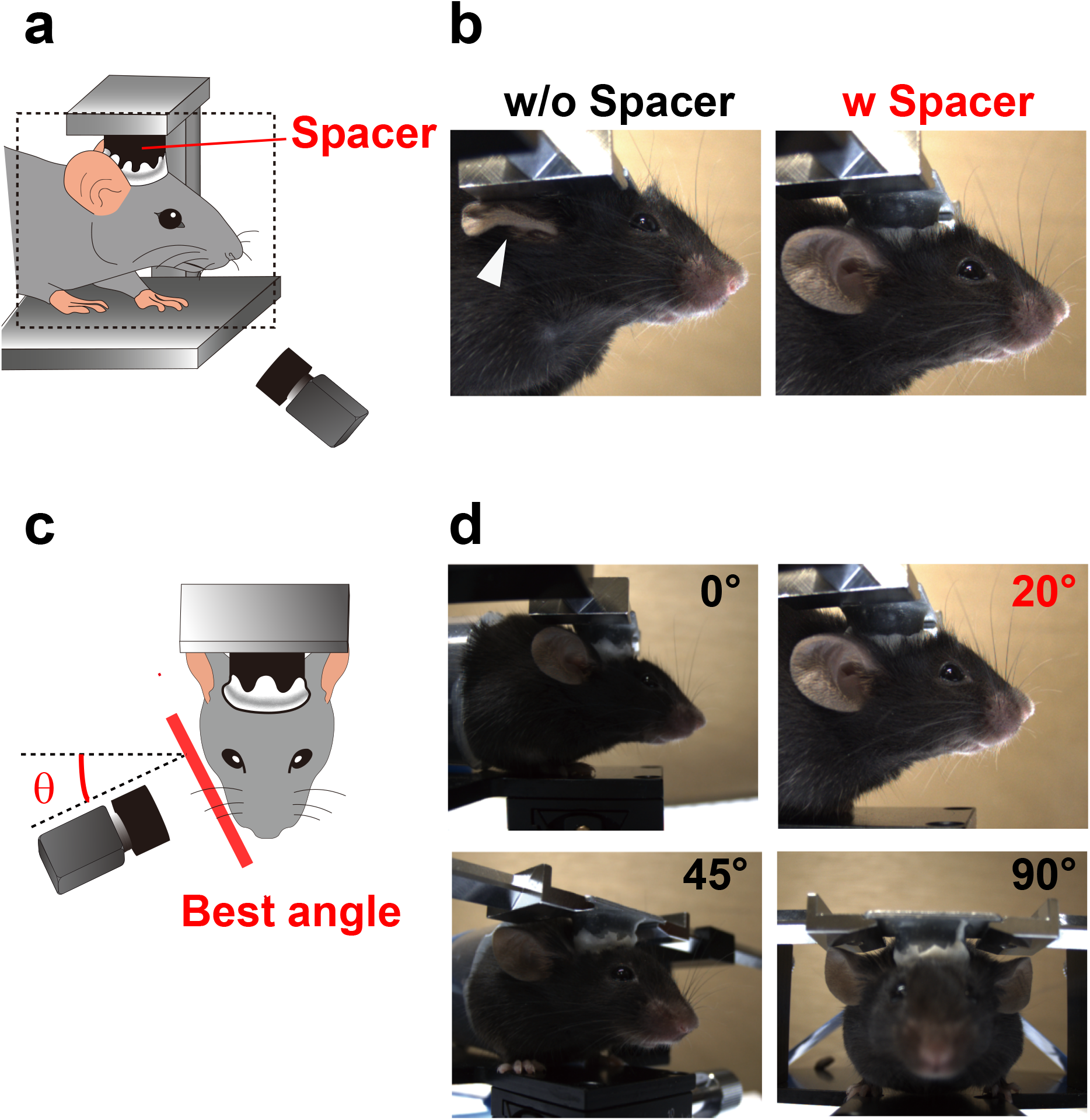
Optimization of facial videography in mice. **a**, Schematic illustrating facial videography in this study. **b**, Facial images captured either without- (left) or with a spacer (right). The spacer prevented ears from interference. **c**, Optimization of the angle for capturing facial expression in mice. **d**, Representative images captured at an angle of either 0°, 20°, 45° or 90° to the anterior-to-posterior axis.

### Facial videography

To acquire the facial recordings, mice were held in a head fixation device (MAG-2, Narishige). The lower half of the body of the mouse was immobilized using a cylindrical tube crafted from a 50 mL conical tube. A uniformly colored wooden board was placed as a background screen. The faces of the mice were illuminated with white-colored light-emitting diodes on the floor, and were photographed from a 20° angle from the sides of the device (Fig. 1c-d). The mice were habituated on the device for a few ten minutes each day at least for a week before the experiments. Facial images were acquired with a color CMOS camera (BFS-U3-23S3-C, FLIR Systems, Wilsonville, OR, USA) at a rate of 5 Hz controlled by a custom-made LabVIEW program (National Instruments, Austin, TX, USA), with minor changes from that used elsewhere (18).

### Stimulation protocols

Mice were subjected to either a painful or tickling stimulus for 5 s, following a 60-s resting period. This procedure was repeated five times, with a 300-s interval between each iteration. The tail of the mouse was pinched with forceps to elicit a painful response, while the abdomen was tickled with a cotton ball to elicit a tickling response. For acquisition of neutral faces, images were taken for 300 s without stimulation after the 60-s resting period. If the head plate was detached from the mouse during the stimulation process, the corresponding video sequence was excluded from the analysis.

### Random image sampling

Images taken during 5 s-period after stimulation were used as sets of painful and tickling facial expressions corresponding to the type of stimulus applied. In addition, images taken for 300 s without any stimulation were also included as neutral images. A total of 3099 painful images, 3999 tickling images, and 73598 neutral images were obtained. For the following analysis, the number of images in each group were adjusted to 3099 by random sampling.

### Image preprocessing

The initial step was to detect a mouse region in an input image (Fig. 3b). We utilized a pretrained model of Detectoron2 for mouse detection, which is a PyTorch-based object detection library developed by Meta AI Research (New York City, NY, USA). To generate masked images, we combined the cropped mouse regions with a uniform background color (RGB = [200,167,122]). The processed images were subsequently resized to 227 × 227 pixels for the following network training.

### Network training

We constructed an AI model for classification of three different facial expressions using the preprocessed dataset. The calculation was conducted using MATLAB software. We performed fine-turning on SqueezeNet v1.1 architecture (16) with initial values of weights that had been trained using ImageNet data (19). Cross-entropy error was employed as the loss function to adjust weights. The learning rate was set at 2 x 10^-4^. The network was trained for 120 iterations with one validation for every three iterations. Images were presented in batches of 512. To avoid overfitting, data augmentation was performed with random flipping along the vertical axis and random translocation of up to 30 pixels in the horizontal and vertical directions.

### Evaluation with Grad-CAM

Heatmaps were generated using Grad-CAM method (17) to visualize the regions of interest by CNN-based machine vision. The calculation was conducted using MATLAB software. In each dataset, the images with top five scores for prediction were overlaid with the heatmaps (Fig. 4).

### Post hoc image analysis

Ten geometrical parameters of facial parts in mice, including mouth opening, jaw angle, ear angle, ear eccentricity, ear perimeter, eye angle, eye eccentricity, eye perimeter, ear to eye angle, and ear to eye distance, were manually analyzed using Fiji (20). We assigned a binary value of 1 for open mouths and 0 for closed mouths. The ten parameters were normalized to z-scores defined by (X-X_mean_)/XSD, where X_mean_ and X_SD_ are the mean, and standard deviation of the parameter values, respectively. Principal components were then calculated from the standardized measurements using scikit-learn for Python3. The principal components were then plotted in 3D coordinates.

## RESULTS

### Optimization of facial image acquisition in mice

Rodents reportedly display various facial movements, including those of the ears, eyes, and mouth, during emotional responses (9–11, 21). Therefore, capturing the key parts clearly is critical to analyze the features during facial expressions. Initially, we attempted to acquire facial movies of head-fixed mice using a commercially available stereotaxic device; however, we discovered that the plate of the holding device often interfered with ear movement (Fig. 1b left). To resolve this issue, we attached a cylindrical spacer between the mouse skull and the plate. With the presence of the spacer, the shape of the ear remained undistorted (Fig. 1a - b). Subsequently, we optimized the angle that enables us to capture the entire facial features in mice (Fig. 1c). Among the tested conditions, image acquisition at a 20° angle relative to the transverse direction of mice heads was found to be the most effective for capturing the key parts in focus (Fig. 1d).

### Acquisition of neutral- and stimulation-evoked facial movies

Using the optimized protocol above, we captured neutral- and stimulation-evoked facial movies in mice. To evoke negative- and positive emotions, we applied “painful” tail pinch and “tickling” abdomen brushing, respectively (10, 11, 22). As shown in the representative images (Fig. 2), mice reacted to both painful and tickling stimuli with robust movement in facial parts such as ear, mouth, and jaw, while they showed no detectable facial responses without stimulation. We repeatedly acquired the data using ten identical animals, and finally prepared the dataset containing 3099 images for each emotional state.

**Fig. 2.**
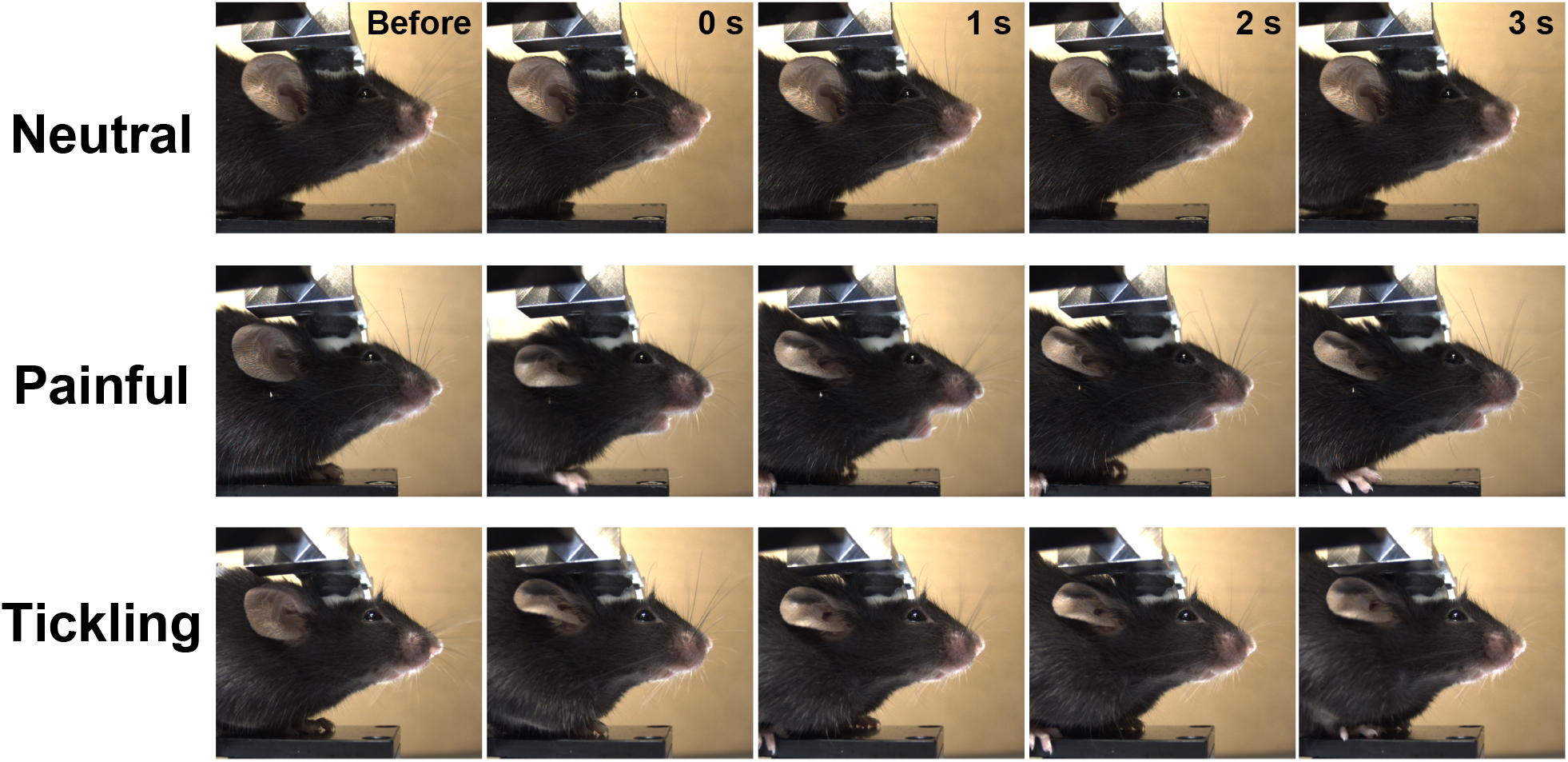
Facial dynamics of a mouse during three emotional states. Time-lapse images for facial dynamics during three emotional states (top: neutral, middle: painful, bottom: tickling). Images were shown at 1-s intervals.

### Framework for our AI-based image classification

Although the patterns of facial movement were subtly distinct among three emotional states (neutral, painful, and tickling), intuitive recognition of the differences was difficult. Therefore, we decided to adopt an AI-based approach for an unbiased image classification. For this purpose, we employed SqueezeNet (16), an eighteen-layer deep CNN model with a high accuracy and fewer parameters, for image classification (Fig. 3a). To remove the information of background objects such as a holding devise in raw images, we conducted masking of the mouse images using a panoptic segmentation algorithm before the trainings in the network model (Fig. 3a-b). By combining the cropped mouse images with the background colors, we generated input images for SqueezeNet. After resizing the images to 227 × 227 pixels, we trained the data in the model. During 120-times iterations of trainings, parameters of learning curves improved well with little to no overfitting (Fig. 3c-d). The model correctly predicted 87.8% of untrained neutral face images (817 out of 930 images), 78.8% of untrained painful face images (733 out of 930 images), and 79.6% of untrained tickling face images (740 out of 930 images), respectively (Fig. 3e). The results demonstrate the validity of our AI model to classify facial expressions during the three emotional states in mice.

**Fig. 3.**
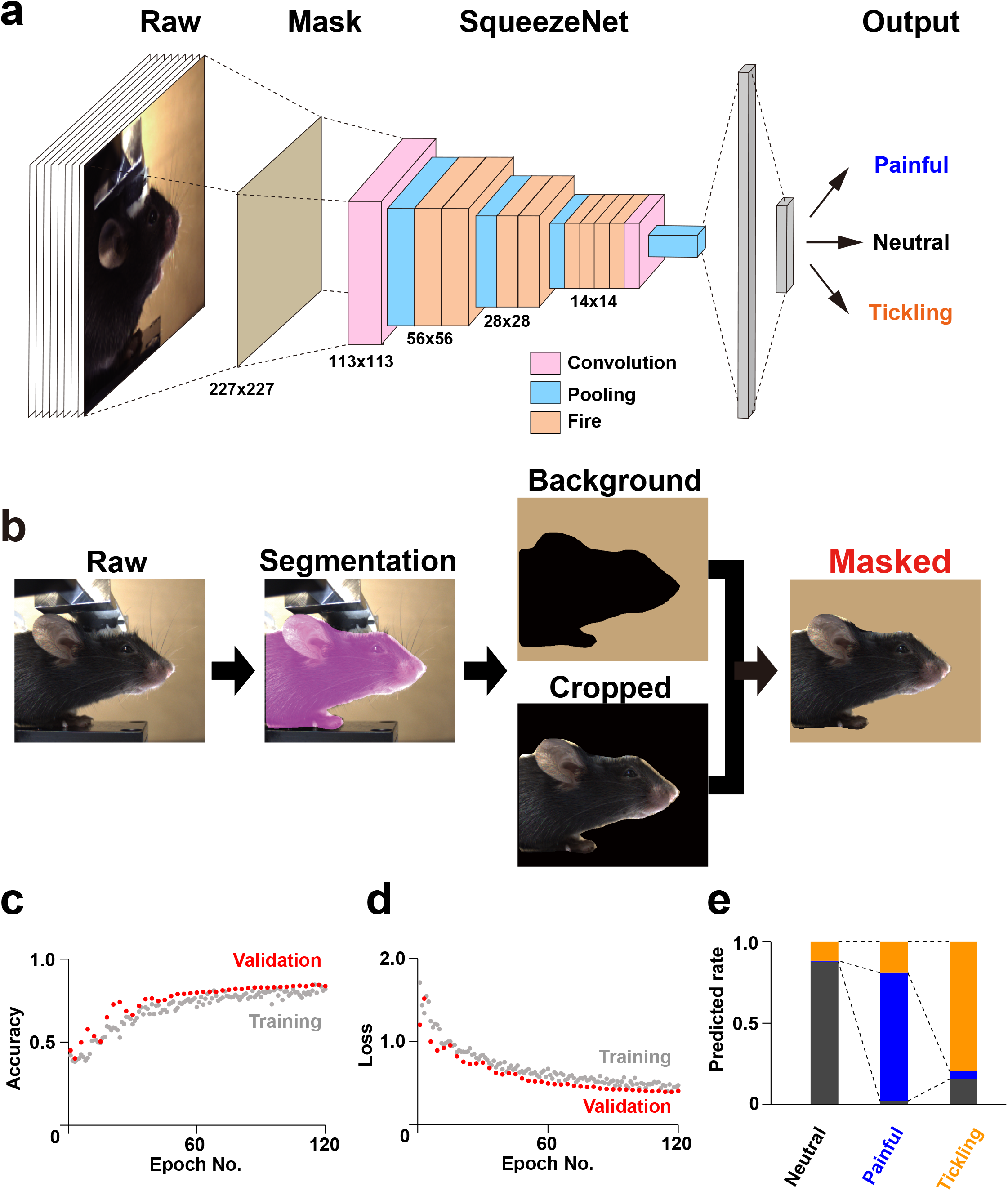
Framework of our AI model for facial recognition in mouse. **a**, Schematic of SqueezeNet-based our AI model for facial recognition in mouse. **b**, Schematic of creating masked images of mice. **c-d**, Learning curves of our model. **e**, Model classification accuracy in validation data.

### Mapping the Facial Features of Each Emotional State

We next analyzed what features our AI model used for the image classification. To delineate the regions of interest of our AI, we employed Grad-CAM (17), which generates a heat map representing the visual rationale for decisions made by CNN-based models. We presented the representative heatmap images with five highest prediction scores for each emotional state (Fig. 4). The results indicate that strong signal appeared around ear, mouth, and jaw. Notably, our model appeared to focus on a flat ear and straight jawline for neutral faces, an open mouth for painful faces, and a crumpled ear and bent jawline for tickling faces, potentially reflecting the dynamics of action units during facial expressions in rodents (9, 10).

**Fig. 4.**
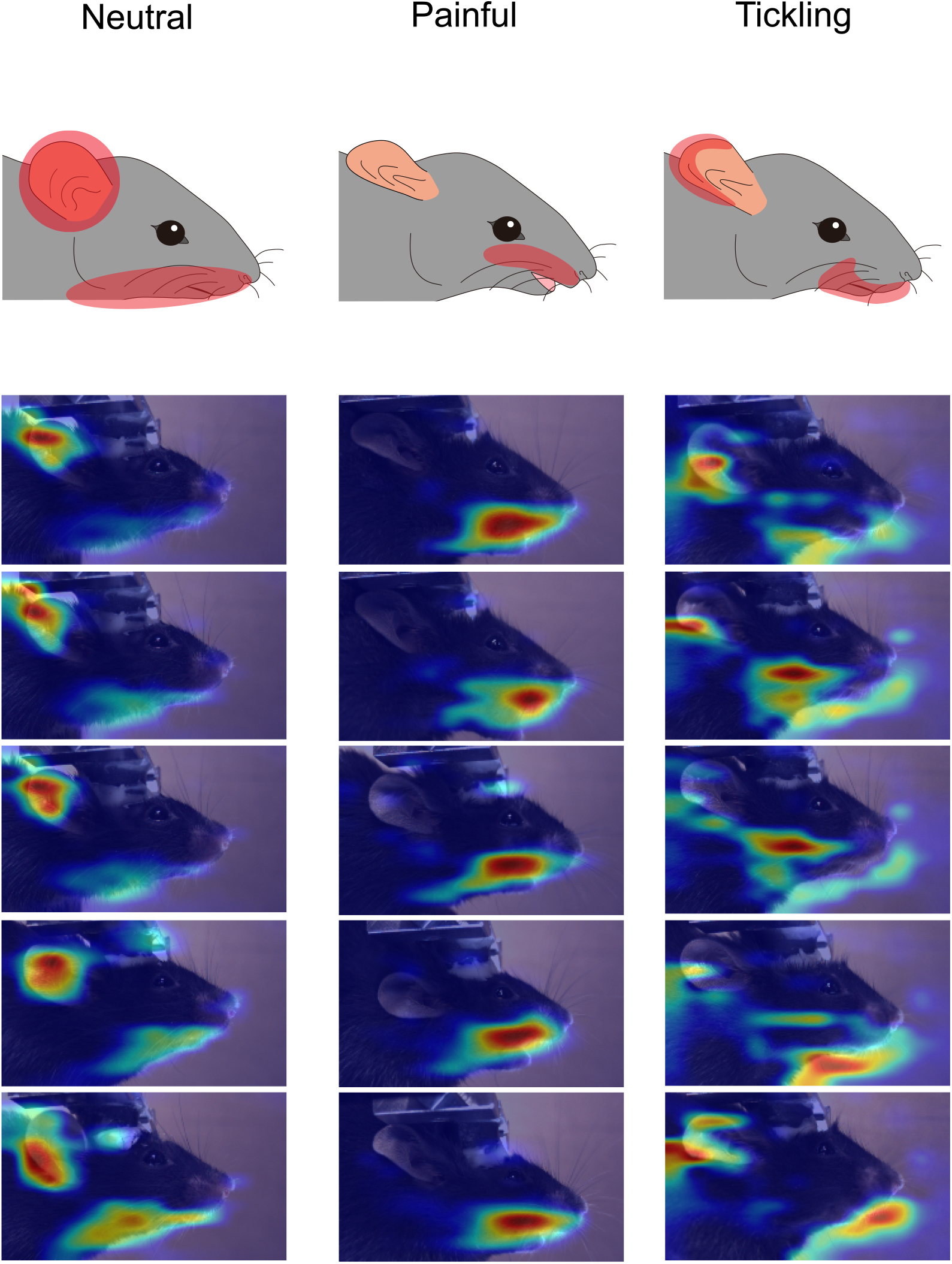
Mapping of the regions where our SqueezeNet based-model focused on. Representative images of Grad-CAM results in each emotional state. Images with top five prediction scores were overlaid with heatmaps created by Grad-CAM analysis.

### Verification of AI Model Interpretations by Post Hoc Image Analysis

To verify the successful prediction of facial features for each emotional state by the Grad-CAM approach, we conducted quantitative analysis on the geometrical parameters for the featured facial parts (ear, mouth, and jaw) as well as eye, a critical parameter for rodent facial expression (9, 10) (Fig. 5a). We quantified ten parameters (mouth opening, jaw angle, ear angle, ear eccentricity, ear perimeter, eye angle, eye eccentricity, eye perimeter, ear to eye angle, ear to eye distance). As evident from the results of Grad-CAM analysis (Fig. 4), open mouth served as a robust hallmark for painful face (Fig. 5b). On the other hand, due to the large variations of most other parameters (Fig. 5c-k), it was somewhat challenging to find out an obvious feature for neutral and tickling faces using a single parameter. Therefore, we conducted dimensionality reduction of the multivariate using principal component analysis. This analysis led to an apparent separation of the three emotional states (Fig. 5l), suggesting that our AI-based model likely discriminated between neutral and tickling faces by leveraging multiple features of facial parts. Together, our post hoc analysis supports the ideas predicted by Grad-CAM analysis.

**Fig. 5.**
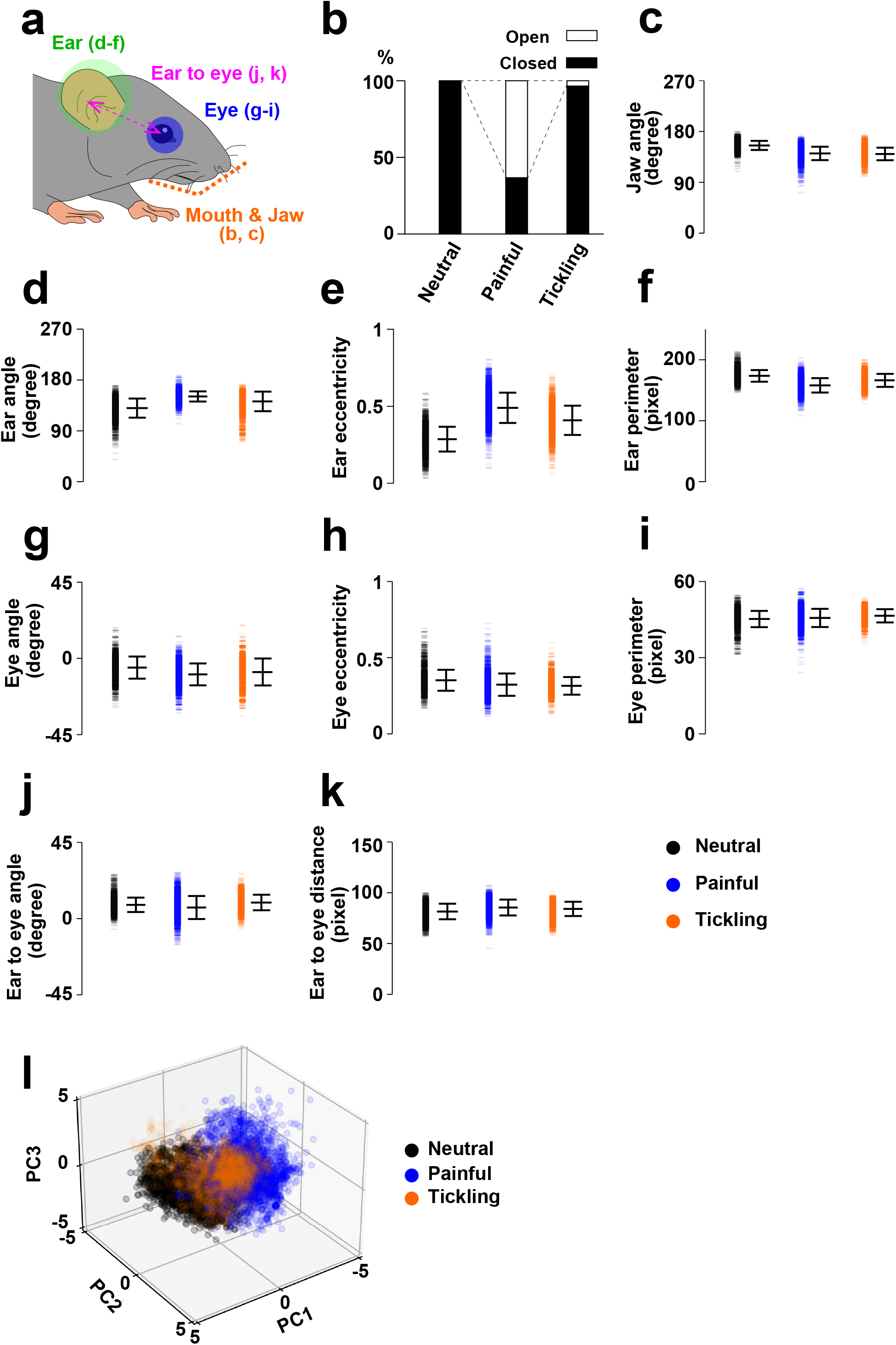
Post-hoc analyses of facial features. **a**, Schematic of analyzed regions in **b-k. b**, Fraction of open or closed mouth. **c-k**, Summary of parameters for the indicated features. Data are shown as mean ± SD. **l**, Scatter plot of the dataset after dimensionality reduction into three principal components.

## DISCUSSION

In this paper, we present a DL-assisted framework for the classification of multiple facial expressions in mice. One of the major benefits of using CNN-based image analysis is that it can automatically learn features in the data, and also can objectively infer the features that contributed to the classification using interpretation techniques such as Grad-CAM. As shown in Fig. 4, our result suggested that our model focused on the facial parts around ear, mouth, and jaw, partially consistent with previous reports (9–11). On the other hand, the regions reported as action units upon emotional events (9, 10), such as eye, were seemingly disregarded by our classifier. Since the facial parts of rodents similarly become active during both painful and tickling emotional events, as described in the previous studies (9, 10) as well as in our study (Fig. 2), the difference in some key features would be too small to characterize the changes among multiple emotional states. Machine vision-based image classification, which takes multidimensional parameters for account, may be superior to manual analyses when detecting such slight differences among multiple facial expressions.

Issues to be addressed are to expand the variation of facial classes. As proposed in a recent study (11), mice likely have several basic emotions similar to Ekman’s basic emotions in humans (2). However, there are a fundamental question: how many patterns of emotion-dependent facial expression do mice possess? Since emotions are thought to originate in the brain, simultaneous recording of facial expression and brain activities, combining our videography with an electrophysiological measurement (23, 24) or fluorescence imaging technique (25), could provide an answer. Expanding our knowledge on the repertoires of facial expression in mice may also require the construction of databases containing a variety of facial expressions in mice, as is widely available for facial expression researches in human (26).

The past decade has witnessed remarkable advancements in artificial intelligence-assisted analysis for big data, owing to the emergence and rapid development of DL-based algorisms (27, 28). DL-assisted tools have made a huge impact on a wide range of research fields, including life and medical sciences. For behavioral researches in mice, such tools have enabled us to automatically capture and track the features of behaving mice, including body postures and vocalization patterns (29–31). Our present work has opened up new avenues for studying the association between facial expressions and emotions in mice. Development of next-generation machine visions that can decode a wider variety of facial expressions more accurately may transform our ability to communicate nonverbally with laboratory animals.

## Author contributions

D.I. conceived the project. H.H. and M.N. supervised the project. Y.T., T.N, and D.I. performed the experiments, analyzed the data, and discussed the results. Y.T. and D.I wrote the manuscript.

## Data Availability

The raw image data used for constructing our network model are available at https://figshare.com/articles/dataset/Dataset_of_Mouse_Facial_Expressions/22083137.

Custom python and MATLAB codes used for data analysis are available at https://github.com/yudaitanaka1026/FacialExpressionAI. Custom LabView codes used for data acquisition are available at https://github.com/daisukeino/FacialExpressionAI.

## REFERENCES

1. Darwin C. The expression of the emotions in man and animals. D Appleton, New York. 1873.

2. Ekman P. Emotion in the human face: Guide-lines for research and an integration of findings. Pergamon Press, New York. 1972.

3. Hart TC, Hart PS. Genetic studies of craniofacial anomalies: clinical implications and applications. Orthod Craniofac Res. 2009;12(3):212–20.

4. Grabowski K, Rynkiewicz A, Lassalle A, Baron-Cohen S, Schuller B, Cummins N, et al. Emotional expression in psychiatric conditions: New technology for clinicians. Psychiatry Clin Neurosci. 2019;73(2):50–62.

5. Gurovich Y, Hanani Y, Bar O, Nadav G, Fleischer N, Gelbman D, et al. Identifying facial phenotypes of genetic disorders using deep learning. Nat Med. 2019;25(1):60–4.

6. Pell MD, Leonard CL. Facial expression decoding in early Parkinson’s disease. Brain Res Cogn Brain Res. 2005;23(2-3):327–40.

7. Parr LA, Waller BM, Fugate J. Emotional communication in primates: implications for neurobiology. Curr Opin Neurobiol. 2005;15(6):716–20.

8. Boneh-Shitrit T, Feighelstein M, Bremhorst A, Amir S, Distelfeld T, Dassa Y, et al. Explainable automated recognition of emotional states from canine facial expressions: the case of positive anticipation and frustration. Sci Rep. 2022;12(1):22611.

9. Langford DJ, Bailey AL, Chanda ML, Clarke SE, Drummond TE, Echols S, et al. Coding of facial expressions of pain in the laboratory mouse. Nat Methods. 2010;7(6):447–9.

10. Finlayson K, Lampe JF, Hintze S, Wurbel H, Melotti L. Facial Indicators of Positive Emotions in Rats. PLoS One. 2016;11(11):e0166446.

11. Dolensek N, Gehrlach DA, Klein AS, Gogolla N. Facial expressions of emotion states and their neuronal correlates in mice. Science. 2020;368(6486):89–94.

12. Li W, Nakano T, Mizutani K, Kawatani M, Matsubara T, Danjo T, et al. Primary motor cortex drives expressive facial movements related to reward processing in mice. bioRxiv. 2022:2022.10.28.514159.

13. Tuttle AH, Molinaro MJ, Jethwa JF, Sotocinal SG, Prieto JC, Styner MA, et al. A deep neural network to assess spontaneous pain from mouse facial expressions. Mol Pain. 2018;14:1744806918763658.

14. Kuo CM, Lai SH, Sarkis M. A Compact Deep Learning Model for Robust Facial Expression Recognition. 2018 IEEE/CVF Conference on Computer Vision and Pattern Recognition Workshops. 2018.

15. Schofield D, Nagrani A, Zisserman A, Hayashi M, Matsuzawa T, Biro D, et al. Chimpanzee face recognition from videos in the wild using deep learning. Sci Adv. 2019;5(9):eaaw0736.

16. Iandola FN, Han S, Moskewicz MW, Ashraf K, Dally WJ, Keutzer K. SqueezeNet: AlexNet-level accuracy with 50x fewer parameters and <0.5MB model size. arXiv. 2017.

17. Selvaraju RR, Cogswell M, Das A, Vedantam R, Parikh D, Batra D. Grad-CAM: Visual Explanations from Deep Networks via Gradient-Based Localization. IEEE International Conference on Computer Vision. 2017:618–26.

18. Ino D, Tanaka Y, Hibino H, Nishiyama M. A fluorescent sensor for real-time measurement of extracellular oxytocin dynamics in the brain. Nat Methods. 2022;19(10):1286–94.

19. Deng J, Dong W, Socher R, Li LJ, Li K, Fei-Fei L. Imagenet: A large-scale hierarchical image database. IEEE conference on computer vision and pattern recognition. 2009:248–55.

20. Schindelin J, Arganda-Carreras I, Frise E, Kaynig V, Longair M, Pietzsch T, et al. Fiji: an open-source platform for biological-image analysis. Nat Methods. 2012;9(7):676–82.

21. Ebbesen CL, Froemke RC. Body language signals for rodent social communication. Curr Opin Neurobiol. 2021;68:91–106.

22. Hinchcliffe JK, Mendl M, Robinson ESJ. Rat 50 kHz calls reflect graded tickling-induced positive emotion. Curr Biol. 2020;30(18):R1034–R5.

23. Jun JJ, Steinmetz NA, Siegle JH, Denman DJ, Bauza M, Barbarits B, et al. Fully integrated silicon probes for high-density recording of neural activity. Nature. 2017;551(7679):232–6.

24. Steinmetz NA, Aydin C, Lebedeva A, Okun M, Pachitariu M, Bauza M, et al. Neuropixels 2.0: A miniaturized high-density probe for stable, long-term brain recordings. Science. 2021;372(6539).

25. Ghosh KK, Burns LD, Cocker ED, Nimmerjahn A, Ziv Y, Gamal AE, et al. Miniaturized integration of a fluorescence microscope. Nat Methods. 2011;8(10):871–8.

26. Yin L, Wei X, Sun Y, Wang J, Rosato MJ. A 3D facial expression database for facial behavior research. 7th International Conference on Automatic Face and Gesture Recognition. 2006.

27. Krizhevsky A, Sutskever I, Hinton GE. ImageNet Classification with Deep Convolutional Neural Networks. Proceedings of the 25th International Conference on Neural Information Processing Systems. 2012;1:1097–105.

28. Alzubaidi L, Zhang J, Humaidi AJ, Al-Dujaili A, Duan Y, Al-Shamma O, et al. Review of deep learning: concepts, CNN architectures, challenges, applications, future directions. J Big Data. 2021;8(1):53.

29. Mathis A, Mamidanna P, Cury KM, Abe T, Murthy VN, Mathis MW, et al. DeepLabCut: markerless pose estimation of user-defined body parts with deep learning. Nat Neurosci. 2018;21(9):1281–9.

30. Sangiamo DT, Warren MR, Neunuebel JP. Ultrasonic signals associated with different types of social behavior of mice. Nat Neurosci. 2020;23(3):411–22.

31. Mathis MW, Mathis A. Deep learning tools for the measurement of animal behavior in neuroscience. Curr Opin Neurobiol. 2020;60:1–11.

